# Biofilm characterization in the maize pathogen, *Fusarium verticillioides*

**DOI:** 10.1101/2022.11.18.517162

**Authors:** Chizné Peremore, Brenda Wingfield, Quentin Santana, Emma T Steenkamp, Thabiso E Motaung

**Author notes:** Correspondence:, Thabiso E Motaung, University of Pretoria, Private Bag X20, Hatfield 0028, Pretoria, South Africa.

## Abstract

Nearly all microbes, including fungal pathogens, form biofilms, which are structured communities of microbial aggregates enclosed in self-produced extracellular polymeric substances (EPS) and attached to a surface. Studying plant-associated fungal biofilms can enhance understanding of fungal biology and knowledge of the links between fungal diseases and plants. However, only a few plant-associated fungi are reported to form biofilms. This study aimed to examine the ability of a mycotoxigenic fungus of maize, *Fusarium verticillioides*, to form biofilms under laboratory conditions. During our investigation, *F. verticillioides* stationary phase cultures incubated in liquid media developed a biofilm-like pellicle with a hyphal assemblage that appears in the form of a cloudy and thin slime material. Under the microscope, the biofilms exhibited a highly heterogeneous architecture made of dense, entangled, and compact hyphae, which were accompanied by a quantifiable EPS and extracellular DNA (eDNA). The biofilm was also found to respond to different abiotic conditions including pH and temperature, suggesting their relevance in a field setting. We further demonstrate the biofilm structural maintenance role of eDNA through treatment with DNase, which was only marginally effective during late biofilm stages, suggesting that it forms complex interactions with the EPS during biofilm maturation. Based on these results, we propose that *F. verticillioides* forms a ‘true’ biofilm that may act as a potential virulence factor.

## 1. Introduction

After rice and wheat, maize (*Zea mays*) is the third most abundant grain crop, feeding millions of people, especially in Sub-Saharan Africa (FAO. 2021). However, its development and yield are threatened by *Fusarium verticillioides*, which systemically colonizes leaves, stems, roots, and kernels. The fungus can therefore induce serious damage that often manifests in *Fusarium* ear and stalk rot (Gai et al., 2018; Oren et al., 2003). These diseases have food safety and security implications due to mycotoxin contamination that is associated with them. Primarily, *F. verticillioides* secretes the mycotoxin fumonisin B_1_, which contaminates symptomatic and asymptomatic maize kernels and stored grains (Munkvold and Desjardins, 1997). The toxicity of this compound is due to the inhibition of ceramide synthase and subsequent toxic intracellular accumulation of sphingosine and other sphingoid bases (Marin et al., 2013), ultimately imposing detrimental health effects on consumer populations (Wild and Gong, 2010). Indeed, studies from around the globe, including some from Africa, Asia, and Latin America, paint a disconcerting picture of how the prevalence of mycotoxins leads to a variety of human pathologies, including oesophageal and liver cancer in adults who consumed contaminated maize (Wild and Gong, 2010).

The mechanisms by which *F. verticillioides* invades maize have been outlined (Gai et al., 2018; Oren et al., 2003), providing important clues on the circumstances leading to infection symptoms and the precise anatomical locations of the maize plant that would probably harbour mycotoxins. Undoubtedly, the production of mycotoxins in *F. verticillioides* has been heavily emphasized over virulence factor production in numerous papers on mycotoxigenic fungi. The current study posits the development of biofilms as a virulence factor is somehow closely related to the accumulation of mycotoxins in fungi. Tell-tale is the biofilm 3D structure and biomass, both of which covered in extracellular polymeric substances (EPS). This structural design gives rise to emergent characteristics that are only seen in the biofilm mode of microbial life, such as surface adhesion-cohesion, spatial organization, physical and social interactions, chemical heterogeneity, and greater tolerance to antimicrobials (Karygianni et al., 2021). In the case of mycotoxigenic fungi, the biofilm EPS may exert a substantial effect on mycotoxin production; it may permit mycotoxins to stably accumulate and persist for longer periods of time due to the fact that that it glues the cells together, in the process creating pockets and channels through which mycotoxins can be concentrated and distributed within a biofilm, respectively. *Aspergillus fumigatus* biofilms, for instance, augment the production of gliotoxin, a sulphur-containing mycotoxin with immunosuppressive properties (Bruns et al., 2010). An interesting area of research will be determining to what extent the components of a biofilm, including the EPS and its associated cell-free components such as the extracellular DNA (eDNA), influence mycotoxin production. Therefore, research on biofilms will provide a fresh perspective on mycotoxin synthesis in economically important fungi.

As biofilms present a cross-sectoral challenge, affecting a wide range of sectors including healthcare, built environment, food and agriculture (Cámara et al., 2022), our lack of understanding of how filamentous fungal biofilms originate, and how they adapt to their microenvironments once developed, will restrict our capacity to identify and counteract their detrimental impacts. This is apparent in the clinical setting, where most clinical infections associated with medical instruments, including indwelling devices (e.g., catheters, pacemakers, dentures, orthopedic prostheses, and heart valves) colonized by biofilms are difficult to treat due to antifungal resistance of these cell community structures (Bryers, 2008; Donlan, 2001; Lindsay and von Holy, 2006). Marine biofilms, on the other hand, which are created by both microorganisms and macroorganisms (such as algae), play a significant role in the environmental effects of biological fouling, also known as marine biofouling, which is the accumulation of undesirable biological matter on artificial submerged surfaces. Ships and underwater surfaces (such as undersea cables and acoustic instruments) are examples of colonized surfaces, and their treatment is difficult due to concurrent and intolerable environmental impacts on non-target species (Callow and Callow, 2011; Bixler and Bushan, 2012). The same type of challenges might apply to agriculture, where some of the essential and most used tools and machinery are contaminated with harmful fungal biofilms that are challenging to remove. When employed across many fields, farming equipment colonized by biofilms has the potential to contaminate unaffected fields with biofilm-derived propagules that may have acquired novel traits within a biofilm, including resistance to fungicides.

Somewhat formal descriptions of biofilms in plant fungal pathogens have started to emerge (Motaung et al., 2020), with the latest being provided for *F. graminearum* (Shay et al., 2022). However, to date, we have no knowledge pertinent to the role of biofilm formation in many disease-causing plant fungi, including key biofilm components such as eDNA and EPS. The current study sought to see if the morphologies of surface-associated development by *F. verticillioides* might be included in current biofilm descriptions. Our research will contribute to a better understanding of how filamentous plant fungal pathogens coordinate survival by forming a community structure.

## 2. Materials and Methods

### 2.1. Fungal strains and maintenance

Strains of *F. verticillioides* were obtained from the Culture collection in the Forestry and Agricultural Biotechnology Institute (FABI), University of Pretoria, following isolation from maize samples in taken from fields in the Eastern Cape province of South Africa. Two strains (CMWF 1196, and CMWF 1197) were screened for their ability to form biofilms in this study. The strains were allowed to grow for two weeks at room temperature by plating on ¼ strength PDA (Potato Dextrose Agar; 10 g of PDA powder and 12 g of Difco agar in 1 L distilled H_2_O). They were then used to develop a biofilm in liquid media as described below.

### 2.2. Screening for biofilm formation

Rapid screening of biofilm formation was performed for all fungal isolates by cutting a block of agar (5 mm x 5 mm) from the sporulating culture and inoculating it into 15 ml of three different growth media. These were: ¼ Potato Dextrose Broth (PDB, Potato Dextrose Broth; 6 g of PDB powder (Difco) in 1 L distilled H_2_O), Roswell Park Memorial Institute-1640 broth (RPMI) and Sabouraud Dextrose Broth (SDB or Sabouraud liquid medium) in 50 ml falcon tubes. After mixing, the inoculate solution was poured into petri dishes (100 mm × 15 mm) and incubated at 25 and 30 °C for 24-72 hrs. They were visually analysed every 24 hrs for biofilm formation which is a hyphal assemblage that appears in the form of a cloudy and thin slime material. Photos were taken with an Epson Perfection V700 Photo scanner.

Biofilm formation for the remaining experiments was conducted using a cell counting method. To do this, the inoculum for counting cells was prepared by adding 2 ml of 1x Phosphate-buffered saline (PBS, 137 mM NaCl, 2.7 mM KCl, 10 mM Na_2_HPO_4_, and 1.8 mM KH_2_PO_4_) onto sporulating culture of *F. verticillioides* growing on ¼ PDA for seven days. The plate was swirled to allow spores to be released into the PBS buffer, and then counted using a haemocytometer placed under a Zeiss Axioskop 2 plus Light Microscope. A volume of 40 μL of fungal cell culture (i.e., PBS spore suspension) was diluted by adding 40 μL of 0.4% Trypan Blue solution (Sigma-Aldrich) into an Eppendorf tube, in order to distinguish between dead and living cells. The concentration of the harvested conidia was adjusted to 1 × 10^5^ conidia/ml in ¼ PDB. Of this 200 μL and left to form a biofilm in 48 wells plates (Corning® Costar® TC-Treated Multiple Well Plates from Sigma-Aldrich) or chamber slides (Lab-Tek® Chamber SlideTM System, 8 Well Permanox® Slide) and incubated for seven days at 25 °C. Once biofilms had matured, the culture was then analysed using different microscopy techniques as explained later.

### 2.3. Analyses of biofilm-like structures

To better describe the biofilms formed by *F. verticillioides*, colony and cell morphology were first analysed on plates containing liquid media (PDB), where the former was examined visually every 24 hrs for 7 days. Following the visual screening, the spores were cultured in ¼ PDB under shaking (180 rpm) (Shake-O-Mat, LABOTEC) and stationary conditions for 72 hrs in chamber slides, harvested and then analysed under a light microscope (Zeiss Axioskop 2 plus Light Microscope (LM)). A scanning electron microscope (SEM) was used to study the ultrastructure of biofilms formed as previously described, in chamber slides or 48 well plates with glass coverslips (LASEC SA). Sample preparation for SEM was performed according to Harding et al. (2010). Samples were examined with Zeiss Ultra PLUS FEG SEM Confocal Laser Scanning Microscopy (CLSM). This was used to analyse of *F. verticillioides* biofilms by forming biofilms in chamber slides at different time points (24 and 7 days) and staining them for 30 mins in a dark room with 100 μl of FUN-1, which fluorescent viability probe for fungi (Chandra et al., 2001). Biofilms were then visualized using Zeiss LM 880 CLSM with excitation wavelength at 488nm and emission at 650nm.

### 2.4. Analysis of biofilm-derived cells

This study considered whether biofilm-derived cells are different from planktonic cells in *F. verticillioides*. To do this, biofilms of *F. verticillioides* were formed as previously mentioned. After 7 days of incubation, cells were extracted from the biofilm by scraping and briefly agitated (30 s) to loosen the cells from the EPS matrix. This suspension was filtered through Mira cloth (Sigma-Aldrich) to separate the cells from the EPS material. In parallel, *F. verticillioides* planktonic cells were scraped from the sporulating cultures on ¼ PDA and added to PBS. The cells were then counted and adjusted to the desired concentration (1 × 10^5^ spres/ml). A volume of 10 μl was aliquoted onto ¼ PDA and incubated at 25 °C for 7 days. The agar plates were then examined to study differences in growth and colony morphology, with three sections of each biofilm-derived and planktonic cells examined with Zeiss Ultra PLUS FEG SEM.

### 2.5. Quantification of biomass, EPS and metabolic activity

Biofilm biomass was determined in relation to the production of EPS and metabolic activity. To measure the biofilm biomass, crystal violet was used as it is known to be a good indicator of the amount of cellular biomass (Cruz et al., 2018). Biomass was then quantified according to the method described by Mello et al. (2016). Biofilms in 96 well plates (Corning® Costar® TC-Treated Multiple Well Plates (Sigma-Aldrich) were fixed for 15 min in 200 μl of 99% methanol. The supernatant was then removed, and the biofilm in microtiter plates was air-dried for 5 min prior to adding 200 μl of 0.3% crystal violet solution (stock solution diluted in PBS; Sigma-Aldrich) to each well. The stained biofilms in micotiter plates were then incubated for 20 min at room temperature prior to being rinsed twice with PBS to remove excess dye. The biomass in each well was then decolourized with 200 μl of 99% ethanol for 5 min. A volume of 100 μl of this solution was then transferred to a new 96-well plate and the absorbance was measured at 590 nm using a microplate reader (SpectraMax ® Paradigm® Multi-Mode Detection Platform).

Non-fixed biofilms were stained for 5 min at room temperature with 20 μl of 0.1% safranin (stock solution diluted in PBS:Sigma-Aldrich). After that, the stained biofilms were rinsed with PBS till the supernatants became transparent. With 200 μl of 99% ethanol, the extracellular matrix was decolorized. A volume of 10 μl of the supernatant was transferred to a fresh 96-well plate, and the absorbance at 530 nm was measured using a microplate reader as previously reported. To measure the metabolic activity of the biofilm, biofilms were formed in 96 well plates as before and incubated at 25 °C. Once biofilms had been formed, their metabolic activity was then quantified using a colorimetric assay, XTT (sodium 3′- [1- (phenylaminocarbonyl)- 3,4-tetrazolium]-bis (4- methoxy6-nitro) benzene sulfonic acid hydrate) (Sigma-Aldrich), according to the manufacturer’s recommended specifications. The activity of fungal mitochondrial dehydrogenase converts XTT tetrazolium salt to XTT formazan resulting in a colour change that can be measured using the microplate reader as previously described, with the absorbance from each well measured at 492 nm.

### 2.6. The impact of abiotic conditions on biofilm growth

The impact of abiotic conditions on biofilm production by *F. verticillioides* was assessed at different pH values (2, 3, 4, 5, 6, 7, 8) and temperatures (10, 15, 20, 25 and 35 °C) in ¼ PDB. The pH was adjusted using HCl and NaOH. To conduct these experiments, standardized spore suspensions were inoculated as previously described and biofilms were allowed to develop under the aforementioned abiotic conditions for 7 days. Quantification of biomass, EPS, and metabolic activity was then performed using the microplate reader as previously described. Following this, EPS and biomass were determined, with EPS expressed per biomass (i.e., EPS/Biomass) and metabolic activity percentages calculated to compare the response of biofilms under the different physical factors.

### 2.7. Effect of DNase treatment on biofilms formation

The eDNA release was measured using a microplate fluorescence assay using a DNA binding dye (SYBR® Green I), as previously described by Rajendran et al. (2014). First, EPS was extracted following a method described by Rajendran et al. (2013). For this study, biofilms were cultivated in ¼ PDB for 7 days at 25 °C before being scraped from the plates using sterile cell scrapers and rinsed with PBS. The EPS was extracted from the disaggregated biofilm using 0.2 M EDTA. Following this the samples were centrifuged at 10,000 x *g* for 30 min, the EDTA supernatant was collected and filtered through a 0.45 m syringe filter (Millipore). SYBR® Green I (Invitrogen), whose binding results in fluorescence that is directly proportional to DNA content, was applied at a 1:4 ratio to biofilm supernatants in a black well microtiter plate (Costar3603; Corning). The levels of eDNA were then quantified using a fluorescence plate reader (SpectraMax^®^ Paradigm^®^ Multi-Mode Detection Platform) at 485 and 525 nm, respectively. The concentration of eDNA in the sample was quantified using the DNA standard curve as previously described by Leggate et al. (2006).

The role of eDNA in *F. verticillioides* biofilm formation was investigated by depletion of eDNA within the biofilm using the hydrolytic enzyme (Rajendran et al., 2013), DNase I from bovine pancreas (Sigma-Aldrich). The DNase I was prepared in 0.15 M NaCl supplemented with 5 mM of MgCl_2_. To assess the effect of DNase I on biofilm formation, biofilms were formed in ¼ PDB as described above and were incubated with DNase I at the concentration of 0.25, 1, and 2 mg/ml, and incubated at 25°C for 72 hrs and 7 days. Untreated controls were included for direct comparison. After each treatment, the biofilms were washed in PBS and their metabolic activity, biomass, and EPS were quantified as mentioned prior.

### 2.8. Statistical analysis

All experiments were performed in duplicate, in two independent experimental sets. The data were expressed as mean ± standard deviation (SD). The results were evaluated using the GraphPad Prism 9 computer program. A *p*-value of 0.05 or below was deemed statistically significant in all analyses (ns= *p* > 0.05; *= *p* ≤ 0.05; **= *p* ≤ 0.01; *** =*p* ≤ 0.001; ****= *p* ≤ 0.0001).

## 3. Results

### 3.1. Colony morphology of *in vitro* biofilm-like structures

The ability of *F. verticillioides* to form biofilm-like structures was observed in both strains examined in this study (Figure 1). Then, using visual inspection of liquid cultures, colonies emerging from a biofilm culture were observed in PDB, SDB and RMPI, incubated at 25 °C from 24 hrs to 7 days (data not shown). The biofilm is normally distinguished from planktonic cells by their dense, highly hydrated clusters of cells enmeshed in a gelatinous matrix (Coraça-Huber et al., 2020; Hurlow and Bowler, 2009, Metcalf and Bowler et al., 2013). Indeed, *F. verticillioides* biofilm-like colonies displayed a dense, thin, and cloudy material (Figure 1). Based on these morphological traits, the biofilm-like formations will be referred to as simply biofilms from this point on.

**Figure 1:**
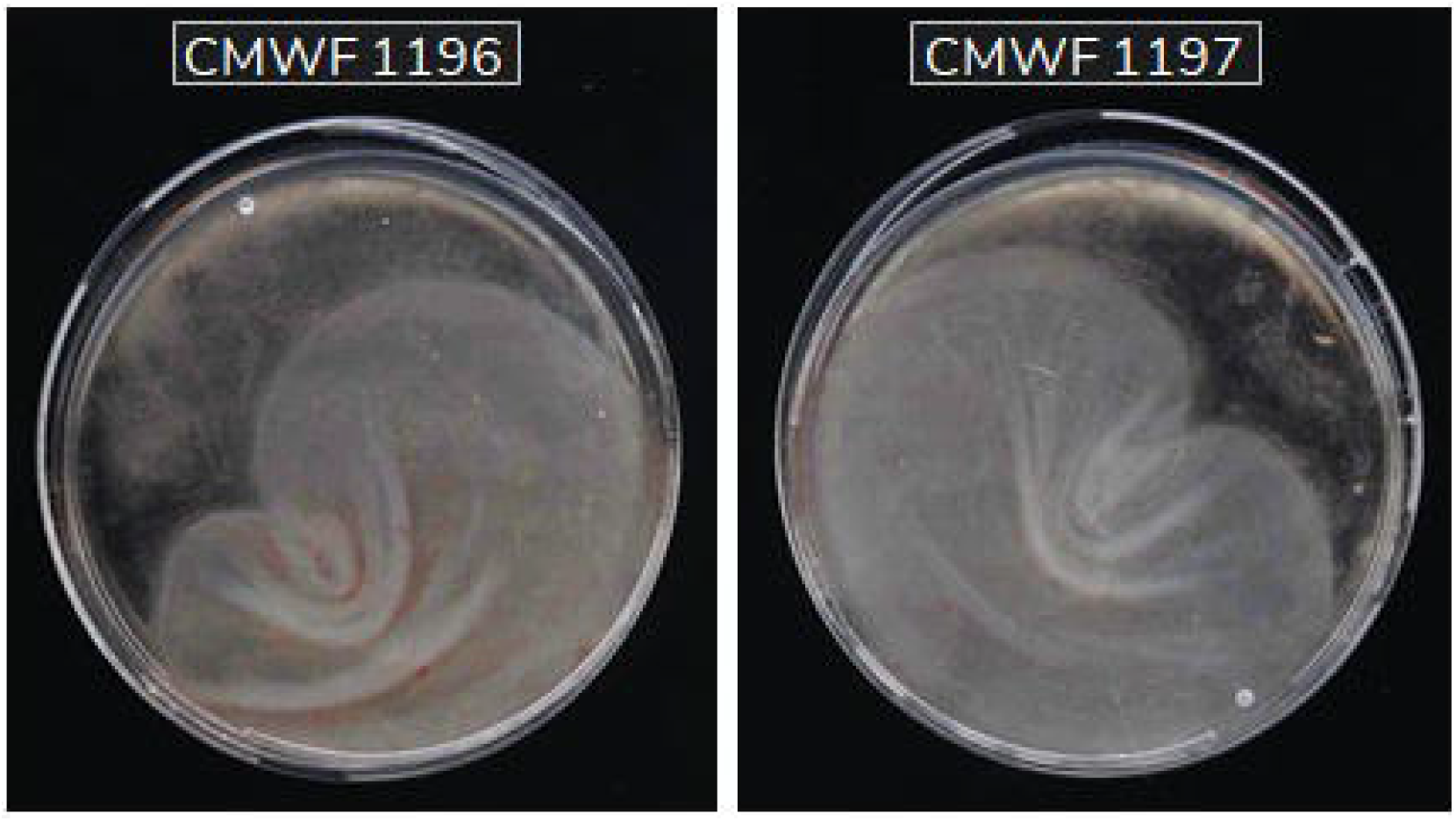
*Fusarium verticillioides* strains that formed biofilm-like cultures in Petri-dishes containing ¼ strength Potato Dextrose Broth. The strains were left to grow, without shaking for 7 days at 25 °C.

### 3.2. *Fusarium verticillioides* biofilm development

Since, morphologically, both the strains appeared to be forming similar biofilms, only one CMW 1196 was then selected and utilized for subsequent studies. It was anticipated that a biofilm would form most effectively under stationary conditions and assumed that the shear stress from shaking would prevent the formation of the EPS matrix, a distinguishing feature of microbial biofilms. Therefore, the cells from *F. verticillioides* CMW 1196 were cultured under both shaking and stationary conditions. Following this, it was found that planktonic cells incubated under shaking conditions did not typically clump together when observed under a light microscope (Figure 2A), but those incubated under stationary conditions developed a community of cells resembling a biofilm (Figure 2B). When these cells were analysed under SEM, little to no EPS formation in cells incubated under shaking conditions was observed (Figure 2C). However, cells that were incubated without shaking formed a visible EPS (Figure 2D), which was not surprising to us given that EPS have been observed often after growth under non-shaking conditions in a variety of biofilm investigations (Cavalheiro and Teixeira, 2018).

**Figure 2:**
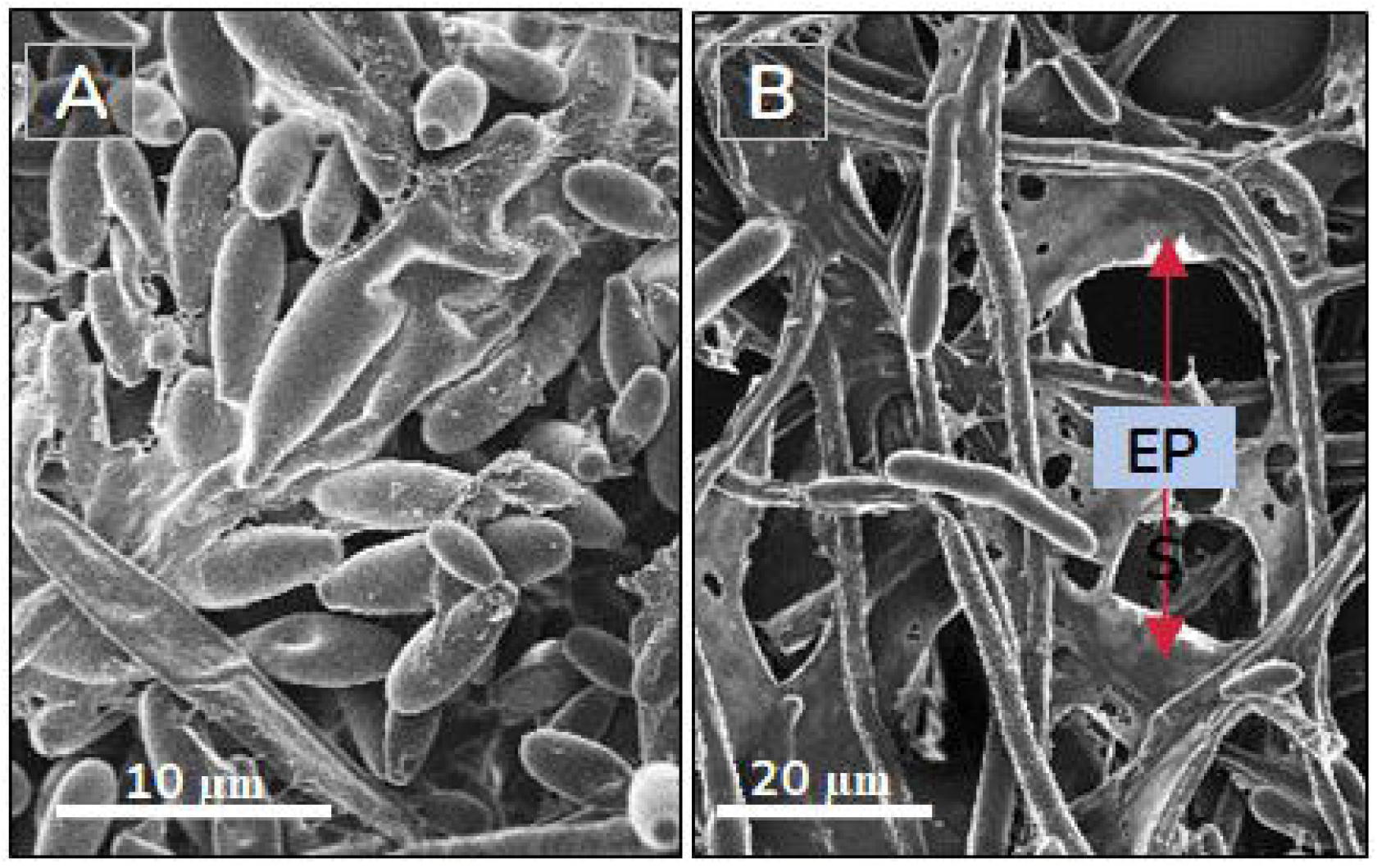
*Fusarium verticillioides* cultured for 7 days under shaking conditions **(A, C)**, remained in the planktonic (free-living) state, and without shaking **(B, D)**, formed biofilms with observable extracellular polymeric substances (EPS, indicated by red arrows in **D**).

### 3.3. *Fusarium verticillioides* biofilms and impact on cells therein

The development of a biofilm in *F. verticillioides* may influence the metabolic status of cells and by extension, phenotype (Ramage et al., 2012). Confirming this were that the spores at the dispersion stage (Figure 3D) appeared to be morphologically distinct from the normal microconidia spores initially used as inoculum to initiate a biofilm (indicated in Figure 3A). Usually, microconidia of *F. verticillioides* are club- or elliptical-shaped or pointed at both ends (Figure 3A). However, the biofilm-derived spores appeared to be more globose/lemon-shaped and slightly larger than typical conidial cells (Figure 3D). Therefore, biofilm-derived cells, as indicated in Figure 3D, may influence phenotypic diversity in *F. verticillioides*. For instance, when these cells were harvested from a biofilm and plated on ¼ strength PDA, they displayed a colony morphology that is different from cells not derived from a biofilm i.e., they formed a colony smaller than that of their planktonic counterpart (Figure 3). This implies that the physiological response of cells contained within a biofilm is greatly influenced by the biofilm ecosystem. However, the morphology of cells derived from a biofilm had no apparent differences when compared to the morphology of normal cells (Supplementary Figure 2), suggesting that the differences between these cells might largely be in their response to environmental signals, as in the case with observations in Figure 3, as opposed to their morphology.

**Figure 3:**
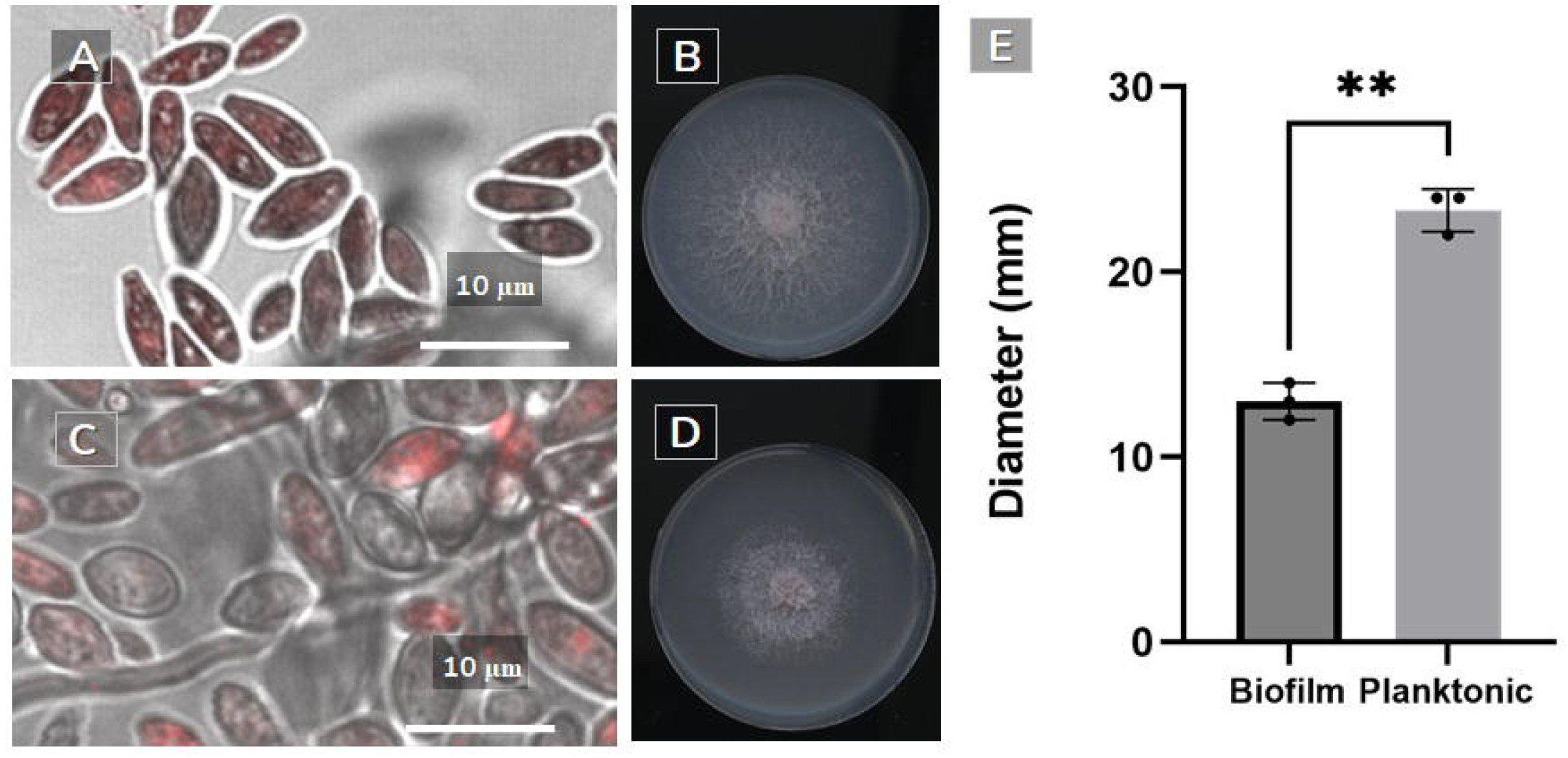
Biofilm-derived and planktonic cultured *Fusarium verticillioides* in chamber slides for (**A**) 24 hrs and (**C**) 7 days. The cells were stained with FUN-1 to determine biofilm developments. *F verticillioides* cells from shaking (**B**) and non-shaking (**D**) cultures established colonies on ¼ PDA. Colonies derived from biofilm cells of asexual cells that were formed were significantly smaller (**=*p* ≤ 0.01) than those derived from planktonic cells (**E**). This is despite the fact that the respective cells were plated at the same concentrations and incubated under the same conditions. Each dot on the bar graphs represents an independent biological replicate. ns= *p* > 0.05; *= *p* ≤ 0.05; **= *p* ≤ 0.01; *** =*p* ≤ 0.001; ****= *p* ≤ 0.0001.

### 3.4 EPS/Biomass and metabolic activity as indicators of biofilm response

The complexity of the biofilm is for the most part brought on by the release of EPS (Ramage et al., 2011). This means that the biofilm may possess the ability to affect the physiology of the cells within it by virtue of holding them in place, thus maintaining the biofilm’s 3D structure while also optimising the exchange of nutrients and genetic material. As previously indicated, the results showed that unlike cells cultured under shaking conditions (Figure 2A, C), the generation of a visible EPS occurs concurrently with the establishment of a mature biofilm (Figure 2B, D). Since establishing that *F. verticillioides* biofilms produce EPS, in this study, we were also interested in how much of this material was being produced during biofilm formation and to what extent metabolically active cells contributed to the total biofilm biomass. For this reason, colorimetric assays were applied, namely crystal violet and XTT, to analyse the biomass and EPS by the absorption of safranin, and to assess the metabolic activity (cell viability) of the biofilm, respectively. In the case of the XTT reduction assay, the production of soluble coloured formazan salts by sessile cells is a direct reflection of cellular metabolic activity. According to the results, an increase in cell mass and EPS production (Figure 4A) seems to be accompanied by an increase in metabolic activity of the cells inside a biofilm (Figure 4B); EPS/Biomass increased significantly from 13% (3 days) to 45% (7 days) (*p* = 0,002), suggesting the more the biofilm matures the more EPS is produced.

**Figure 4:**
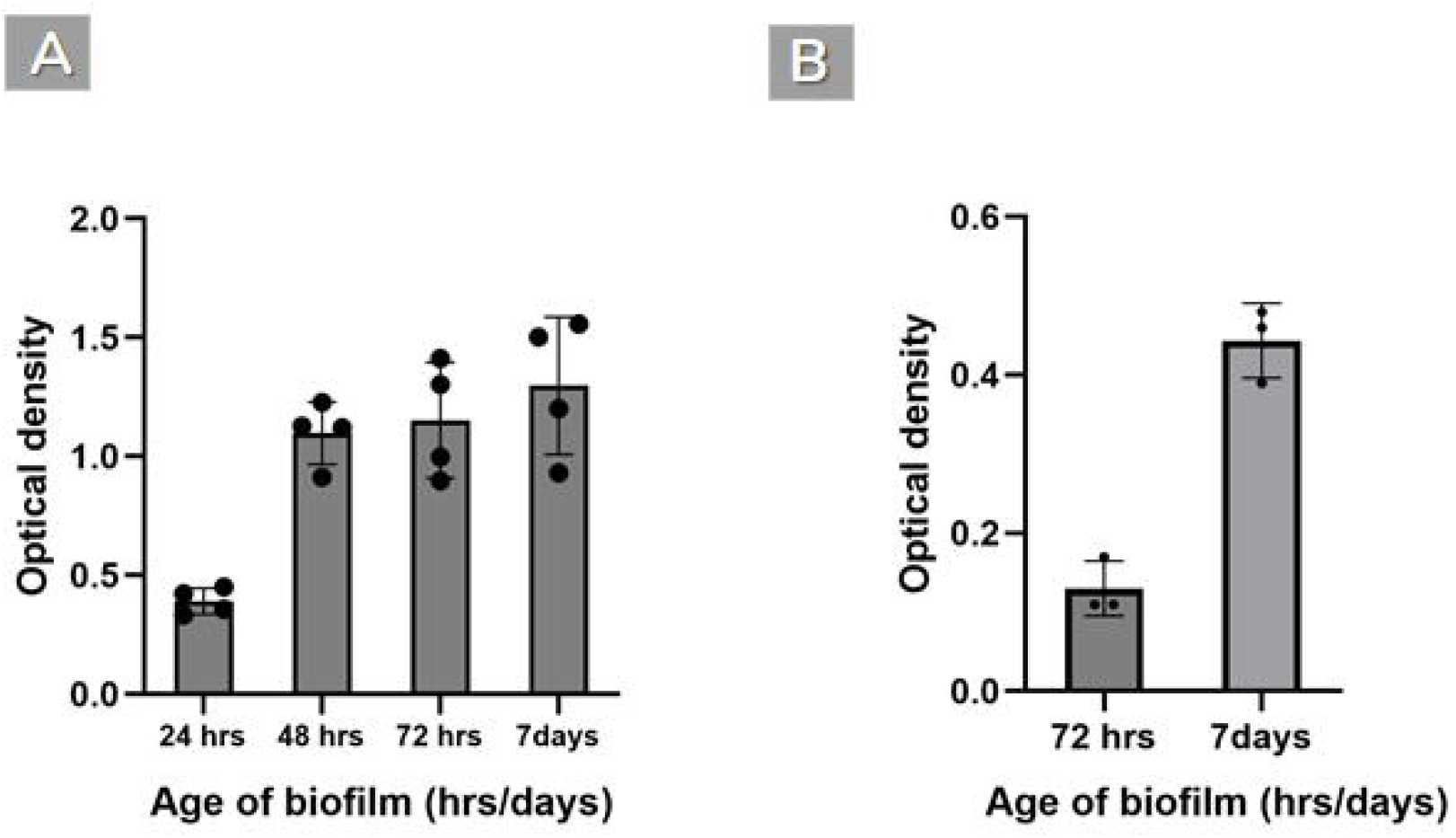
*Fusarium verticillioides* biofilm formation was assessed based on (**A**) metabolic activity, evaluated byXTT reduction assay and (**B**) biomass and extracellular polymeric substances (expressed as EPS/Biomass) evaluated using crystal violet (OD_590nm_) and safranin (OD_530nm_), respectively (*p*-value=0.0013). An increase in cell mass and EPS production is observed. Each dot on the bar graphs represents an independent biological replicate. ns= *p* > 0.05; *= *p* ≤ 0.05; **= *p* ≤ 0.01; *** =*p*≤ 0.001;****= *p*≤ 0.0001.

### 3.5 Biofilm formation in response to abiotic factors

Having shown that *F. verticillioides* asexual cells could develop into a biofilm, this study then investigated how the biofilm will react to different environmental conditions, namely different temperature and pH conditions. As shown in Figure 5A, EPS/Biomass was highest at pH 5, suggesting that *F. verticillioides* may prefer pH 5 for biofilm formation. The media that was used in the initial experiments (¼ PDB) has a pH of around 5, and it was in this medium that all the stages of a biofilm were observed (Figure 2). The EPS and metabolic activity were essentially the same at acidic pH levels (i.e., 2, 3, and 4, below the optimal pH of 5), whereas at pH levels higher than the optimal (i.e., pH 6, 7, and 8, higher than pH 5), the EPS was produced at lower levels than at pH 5 and essentially stayed at these levels. From the optimal pH (pH 5) point of view, *F. verticillioides* biofilms seem to produce significantly more EPS/Biomass than at pH 6 and pH 8 (*p* ≤ 0.05). However, this biofilm had a significantly lower metabolic activity at a range of pHs from 2-8 (2, 3, 4, 6, 7, and 8) (*p* ≤ 0.001). This suggests that, although pH 5 permits better biofilm formation, the biofilm response to pH is versatile, spanning a range of pH conditions, which could influence the adaptability of this pathogen to a range of field conditions.

**Figure 5:**
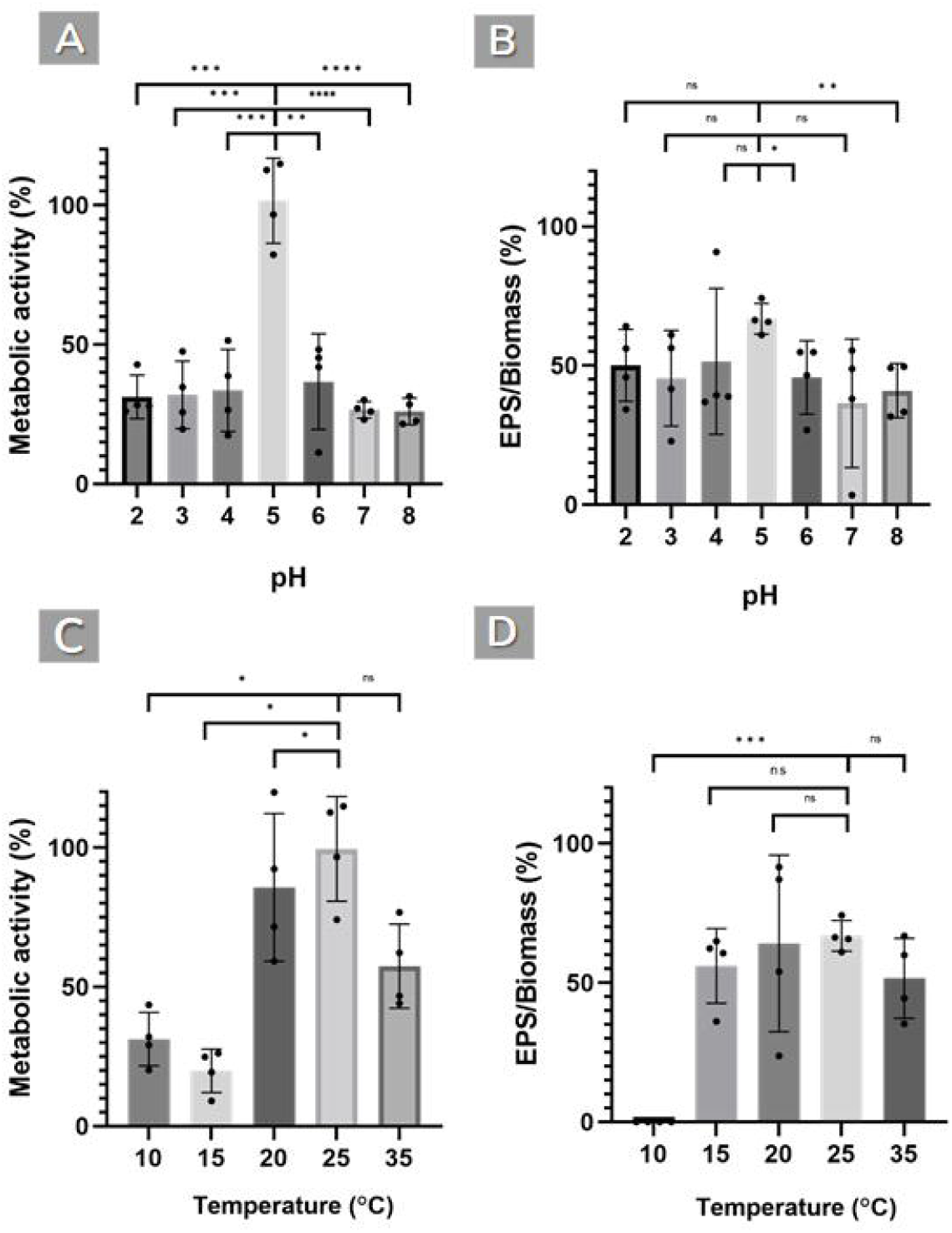
*Fusarium verticillioides* biofilm formation assessed at various pH and temperature conditions by measuring (**A, C**) metabolic activity (XTT reduction assay (OD_475nm;_ p value<0.05) (expressed as metabolic activity percentage)), and (**B, D**) biomass and extracellular polymeric substances (expressed as EPS/Biomass percentage), evaluated using crystal violet (OD_590nm_)and safranin (OD_530nm_), respectively (*p*value<0.05). This data shows that pH 5 permits better biofilm formation, and the biofilm response to pH is versatile, spanning a range of pH conditions. The optimum temperatures evaluated for metabolic activity were 20 and 25°C. It also generated biofilms at 10,15, and 35°C, but these were less robust and had lower metabolic activity. Different temperatures do not appear to have a significant impact on the EPS/Biomass %, except for 10°C, when the biomass could not be quantified. Each dot on the bar graphs represents an independent biological replicate. ns= *p* > 0.05; *= *p* ≤ 0.05; **= *p* ≤ 0.01; *** =*p* ≤ 0.001; ****= *p* ≤ 0.0001.

Figure 5C and D depict the influence of temperature on the production of *F. verticillioides* biofilms. The best temperature tested for metabolic activity is 25 °C, and the fungus displayed similar metabolic activities and formed robust biofilms at 20 and 25 °C with no significant differences. The fungus also formed biofilms at 10, 15 and 35 °C, but these were not as robust as 20 and 25 °C and had lower metabolic activity. Different temperatures do not seem to significantly affect the EPS/Biomass percentage except for 10 °C where the biomass was beyond detection.

### 3.6 The structural maintenance role of extracellular DNA

The biofilm was treated with DNase I to show how eDNA maintains the structure of the biofilm. The DNAse caused the collapse of biofilm formation during early stages of development i.e., at 72 hrs (Figure 6A). Also, the structural integrity of the biofilm was revealed to be significantly impacted by the addition of DNase I in a concentration-dependent manner. Unfortunately, the EPS/Biomass percentages could not be determined during the early biofilm maturation phase (72 hrs) as the biofilm was too weak to perform the relevant assays. In comparison to the biofilm growth control at 7 days, the application of 0.25 and 1 mg/ml, DNase I slightly reduced EPS/Biomass by 17% (*p* > 0.05) while 2 mg/ml significantly reduced it by 40% (*p* = 0.0095) (Figure 6B). Interestingly, although DNase I had an impact on the EPS/Biomass, the metabolic activity of the biofilm was not drastically affected (Data not shown), which could show that eDNA has more impact on the structure of *F. verticillioides* biofilm than on cellular activity, as has been demonstrated in other fungi

**Figure 6:**
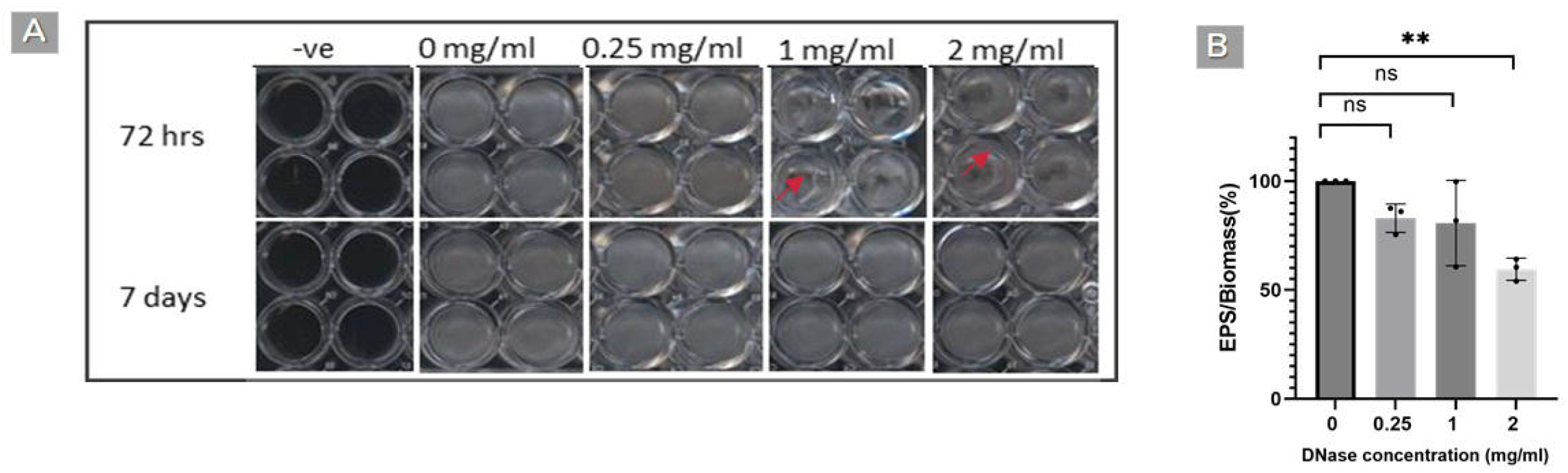
*Fusarium verticillioides* biofilm response to DNase treatment. (**A**) The response of a 72hr-old biofilm to DNase at different concentrations (0.25,1 and 2 mg/ml). (**B**) The response of biofilms measured in biomass and extracellular polymeric substances (expressed as EPS/Biomass percentage). This data shows that DNase I caused the collapse of biofilm formation at the early stages of growth, i.e., at 72 hrs, and the structural integrity of the biofilm was found to be strongly influenced by DNase I in a concentration-dependent manner. Each dot on the bar graphs represents an independent biological replicate. ns= *p* > 0.05; *= *p* < 0.05; **= *p* ≤ 0.01; *** =*p* ≤ 0.001; ****= *p* ≤ 0.0001.

## 4. Discussion

Members of the genus *Fusarium* cause economically important and hard to control diseases including cankers, crown rot, head blight, scabs, and wilts. Although many of these diseases likely have strong links with biofilm formation (Harding et al., 2010; Harding and Daniels, 2017; Motaung et al., 2020) given that microbes predominantly exist in a state of a biofilm in their natural environments, biofilm formation has been formally described in just a few *Fusarium* species including *F. oxysporum* f. sp. *cucumerinum* and *F. graminearum* (Li et al., 2014; Shay et al., 2022). Therefore, for many fungal pathogens of plants, including those belonging to *Fusarium*, it is unclear how biofilms form, let along how they impact infections and disease outcomes.

Seven years ago, Miguel and his colleagues saw what looked like *F. verticillioides* biofilms, in which the mycelium was structured in an extracellular material around the hyphae (Miguel et al., 2015). These researchers also discovered a flocculent substance over the cells or small fibrils of hyphae connecting to one another, like a biofilm. To the best of my knowledge, this is the only time that evidence of a biofilm-like structure for *F. verticillioides* has been reported *in vitro*. There is no formal description of biofilms by this fungal pathogen. In the current study, we addressed this knowledge deficit by describing how *F. verticillioides* forms biofilms under *in vitro* conditions.

Information that describes biofilm colony morphologies in stationary liquid cultures is generally lacking in fungal plant pathogen biofilm studies. The filamentous fungus *F. graminearum* has recently been described as being able to form biofilm colonies at the air/liquid interface that are growing as pellicle–floating masses of cells that cling to each other and move as a unit (Shay et al., 2022). In Figure 1 of this study, it is shown that floating masses for *F. verticillioides* can be distinguished from free-living (planktonic) cells by forming colonies displaying a dense, thin, and cloudy material. The results of this study are consistent with several other studies also reporting similar features in plant and animal tissues, as well as surgically implanted equipment such as catheters and pacemakers colonized by fungal and bacterial biofilms (Coraça-Huber et al., 2020; Hurlow and Bowler, 2009; Santos et al., 2011; Trautner and Darouiche, 2004).

In the current study, it was also observed that biofilms in *F. verticillioides* also developed most efficiently under stationary conditions, while shear stress from shaking conditions prevented proper biofilm formation. Cells incubated in the stationary conditions without shaking had hyphal cells tangled with EPS that appeared to behave like a matrix binding the hyphae together, and cells cultured under agitated conditions seldomly clumped together or not at all. These findings are similar to those of Hawser et al. (1998), who demonstrated that only a few cells are visible on the surface of cultures when they are shaking, in contrast to stationary conditions, whereby cultures are characterized by dense networks of hyphae. This might be due to the severe shaking influencing cell architecture, matrix deposition, and biofilm formation (Soll and Daniel, 2016). In *C. albicans*, shaking at a speed of 60 rpm prevents biofilm growth, with biofilms exposed to shear stress being thinner than those exposed to non-shaking conditions (Cavalheiro and Teixeira, 2018). Hawker et al. (1998) also found that lower speeds result in the production of biofilms with no extracellular matrix but only hyphae, while shaking at higher-speed results in a biofilm which consists of a few cells on the surface. In this study, it was observed that some EPS material is present in planktonic cultures (Figure 2C); however, this was not as abundant as in biofilm cells (Figure 2D), suggesting that under conditions causing agitation of the fungal cells, the cells struggle to produce the EPS matrix.

In light of the above, the findings presented here align with the biofilm concept previously proposed (Córdova-Alcántara et al 2019; Harding et al. 2009, 2010; Motaung et al., 2020). The biofilms of *F. verticillioides* seem to develop through spore adhesion, microcolony formation, maturation, and dispersion. Although many fungal and bacterial species have recorded comparable developmental stages for biofilm formation, filamentous fungal biofilm formation seems to differ from strain to strain (Li et al., 2014; Mowat et al., 2007; Ramage et al., 2011,2012). In contrast to unicellular life forms such as yeast and bacteria, most fungi contain many planktonic forms that can disperse and continue the cycle (e.g., sporangia asexual spores, sexual spores, and hyphal fragments), and these dispersive forms most usually float in air rather than water (Harding et al., 2009). In this study, the dispersal phase of biofilms leads to a substantial number of free-living cells in the form of conidia, but these cells appeared to be morphologically distinct from the normal microconidial spores initially used as inoculum to initiate a biofilm (Supplementary Figure 1). This suggests that the dispersed biofilm cells differ from normal microconidia. Similar findings were reported in a study on *Bacillus cereus* where the cells in the biofilm have different cell-surface characteristics than their planktonic counterparts. For example, the structure of a polysaccharide linked to peptidoglycan in *B. cereus* has been discovered to change during biofilm development (Majed et al., 2016). In addition, according to Boles et al. (2004), the short-term development of *P. aeruginosa* in biofilms causes considerable genetic diversity in the resident bacteria. They also discovered that genetic diversity forms bacterial subpopulations in biofilms with specialized functions and that functional diversity improves the biofilm community’s capacity to endure physiological stress. Equivalent diversity has not been shown in fungal biofilms and should be investigated further.

The biofilm detachment phase is important in disease development, and there has been growing evidence that specific techniques of dispersion can result in releasing biofilm cells that are more pathogenic than their planktonic counterpart (Beitelshees et al. 2018). The dispersed biofilm-derived conidia may be critical to the pathogen’s invasive nature during fungal infections. In other studies, biofilm-derived cells have frequently been shown to be resistant to antifungals and stress (Guilhen et al., 2017). Therefore, *F. verticillioides* biofilm-derived conidia need to be further investigated for agriculturally relevant traits including heightened antifungal resistance against fungicides currently employed in agriculture, which could make the disease caused by this fungus challenging to treat.

Traditional methods for probing biofilm biomass, extracellular matrix, and metabolic were used in this study. In *A. fumigatus* biofilm have been previously linked with metabolism, biomass, and hyphal growth (Mowat et al. 2007). It was also shown that metabolic activity and biomass increased during the first 24 hours of biofilm formation and subsequently reaching a plateau of development (Mowat et al. 2007). Mello et al. (2016) focused on the assessment of biofilm formation by *Scedosporium* species and reported similar outcomes in some species in terms of biomass, metabolic activity, and EPS production. The production of the matrix has a high energetic cost, which may be evolutionary justified given the matrix’s structural and physicochemical significance in the growth and operation of the biofilm, without which the beneficial emergent properties of biofilms would not be possible (Flemming et al., 2016; Saville et al., 2011). This goes to show that the formation of biofilms is closely linked to the formation of the matrix, the bulk of which is extracellular material (Li et al., 2014).

As biofilms grow, metabolic activity and EPS production also rise, suggesting that they may become more resistant to abiotic stress (Motaung et al., 2020). In the current study, it was found that *F. verticillioides* developed biofilms under a range of pH and temperature conditions, and a similar trend was previously reported (Li et al., 2014). These results show that *F. verticillioides* biofilms are produced under different conditions and have an optimum pH of 5 and an optimum temperature of 25 °C. Under field conditions it has been reported that the optimum temperature for spore development is 27 °C a temperature at which biofilm development could occur. Evaluation of biofilm formation under different pH conditions showed that biofilms were much more robust in somewhat acidic (pH 4-5) conditions than in acidic (pH 2, 3), neutral (pH 7) or alkaline (pH 8) environments. These findings are consistent with other reports (Cornet and Gaillardin, 2014) since the pH range for fungal development is fairly broad, ranging from pH 3.0 to more than pH 8.0, with the optimum at pH 5.0 assuming nutritional needs are met. The capacity to develop a biofilm under varying physical conditions may provide the fungus with survival benefits in inhospitable environments.

Unlike other components of the biofilm EPS matrix, the eDNA has by far attracted the most attention and is considered a useful tool to study the biology of biofilms. Many studies have uncovered that eDNA plays many important roles in bacteria such as in biofilm structural maintenance, assisted by the action of DNA binding proteins (Buzzo et al., 2021; Kavanaugh et al., 2019; Whitchurch et al., 2002), antimicrobial resistance (Okshevsky and Meyer, 2015; Rajendran et al., 2013), and acting as a reservoir for the interexchange of genes through natural transformation (Merod and Wuertz, 2014). Furthermore, eDNA assumes more unusual roles, including acting as a source of energy and nutrients (e.g., carbon, nitrogen and phosphorus) (Ibáñez de Aldecoa et al., 2017; Mulcahy et al., 2010; Pinchuk et al., 2008), and forming higher-order conformations (e.g., G-quadruplex DNA) that further strengthen the biofilm through extracellular EPS-eDNA networks (Seviour et al., 2021). Given these roles, eDNA is an attractive target of antimicrobial drugs to manage biofilm-related infections. However, there are only a few studies exploring the existence and function of eDNA in filamentous fungi, with no studies conducted in plant pathogenic fungi. In the current study, it was demonstrated that DNase I has an impact on the biofilm stability. In a similar study using *A. fumigatus*, DNase I was effective at all concentrations (0.25, 1, 4 mg/ml) (Rajendran et al., 2013, 2014), but the maximal effect was observed with 4 mg/ml of DNase I (*p* value < 0.001) which is similar to the observations in this study. The discovery that eDNA contributes significantly to the biofilm EPS in both bacteria and fungi suggests that this may be a conserved and possibly active microbial biofilm process. Nucleic acids are incorporated early in the development of the EPS matrix in *F. graminearum*, and they similarly appear to function as a scaffold, likely regulating the entire matrix structure of the biofilm (Shay et al., 2022). The finding presented in the current study imply that eDNA plays a significant structural maintenance role during biofilm development in a filamentous plant fungal pathogen and may contribute to the severity of disease.

## 5. Conclusions

The study showed that the maize fungal pathogen, *F. verticillioides*, can develop a biofilm under laboratory conditions. Even though the description described here of fungal biofilms may have limitations when the natural environment is considered, at the very least *F. verticillioides* biofilm development seems to follow a typical model that entails attachment, colonization, development (including EPS synthesis), maturity, and dissociation from EPS, with the last phase producing conidia to restart the biofilm cycle. However, in the natural environment, plant stem and leaf surfaces can be sparsely or densely colonized by diverse fungal biofilms and are likely more complex than conditions used in this study. Laboratory-grown biofilms are a simple surface-covering, frequently exhibiting confluent and compact uniformity that is consistent with the original definition of biofilms (Motaung et al., 2020). Documenting biofilm formation in the natural environment by analysing heavily infected maize plants and a population of field strains, as opposed to a few ones, is needed to better understand what is happening in field conditions. By better understanding the complexity of plant-associated fungal biofilms and their phenotypic traits, it would be possible to develop novel antifungal drug targets and treatment alternatives to decrease the prevalence of fungal infections. This study thus establishes a baseline with regards to *F. verticillioides* biofilms, showing its intricate structure and response to the environment. The findings presented further showed that eDNA degradation causes biofilms to collapse, suggesting it is a candidate for antifungal development. These findings represent the first investigation on eDNA analysis in a plant fungal pathogen. Taken together, *F. verticillioides* ability to form biofilms may give it an ecological edge in its battle to keep its place as a commensal and pathogen of maize. The biofilm might enable this fungus to evade host immunity, withstand antifungal treatment and competition from other microbes. The current study, together with earlier studies, therefore, will deepen the understanding of the relationship between disease outcomes and biotic interactions in *F. verticillioides*.

## 7. Acknowledgements

Funding from the National Research Foundation (South Africa) through the Thuthuka funding instrument (grant no. 129580) is acknowledged. Any conclusions and/ or recommendations made here are solely the responsibility of the author(s), and the NRF accepts no accountability in this regard.

