## Supplementary figures and images for "Biofilm characterization in the maize pathogen, *Fusarium verticillioides*"

### Figure S1

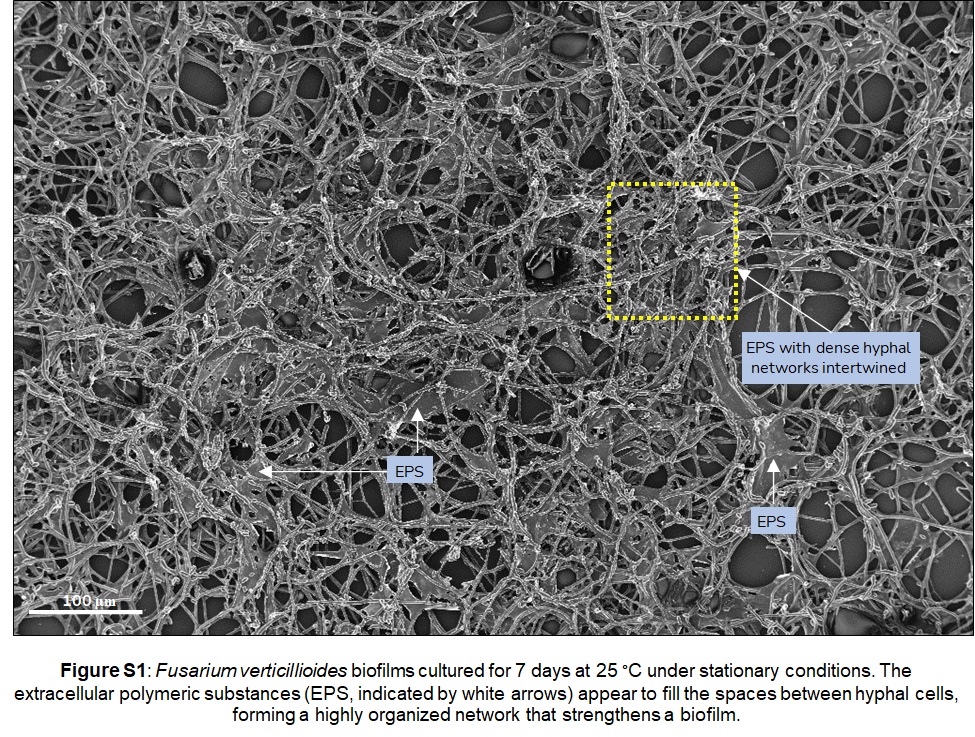

### Figure S2

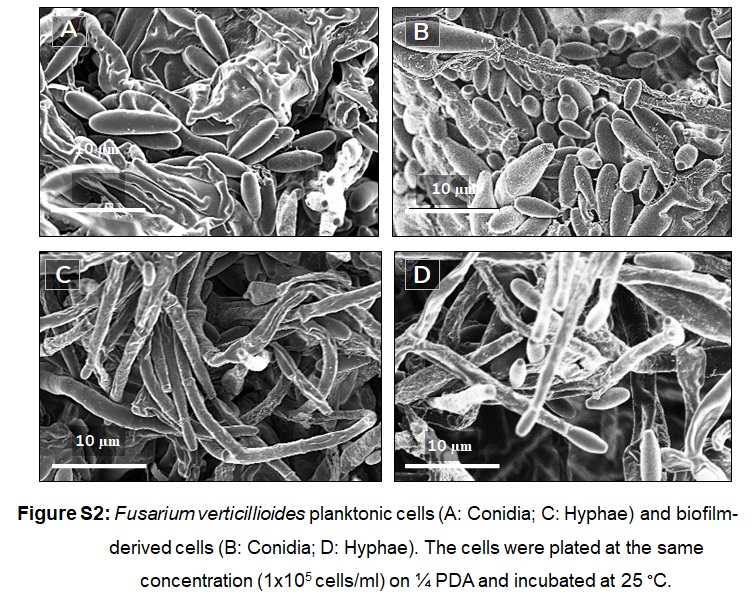
